# A Distinct Neural Code Supports Prospection of Future Probabilities During Instrumental Information-Seeking

**DOI:** 10.1101/2023.11.27.568849

**Authors:** Nicholas M. Singletary, Guillermo Horga, Jacqueline Gottlieb

## Abstract

To make adaptive decisions, we must actively demand information, but relatively little is known about the mechanisms of active information gathering. An open question is how the brain estimates expected information gains (EIG) when comparing the current decision uncertainty with the uncertainty that is expected after gathering information. We examined this question using fMRI in a task in which people placed bids to obtain information in conditions that varied independently by prior decision uncertainty, information diagnosticity, and the penalty for an erroneous choice. Consistent with value of information theory, bids were sensitive to EIG and its components of prior certainty and expected posterior certainty. Expected posterior certainty was decoded above chance from multivoxel activation patterns in the posterior parietal and extrastriate cortices. This representation was independent of instrumental rewards and overlapped with distinct representations of EIG and prior certainty. Thus, posterior parietal and extrastriate cortices are candidates for mediating the prospection of posterior probabilities as a key step to estimate EIG during active information gathering.

## Introduction

In systems neuroscience, decision-making is modeled as a choice between alternative options based on the decision makers’ preferences, goals, and knowledge regarding the choice. Traditional decision-making research has typically applied this framework to simple conditions, in which decision-makers are assumed to possess the information relevant to their choice; for example, a participant in a decision-making experiment is typically given a set of relevant stimuli and asked to make a decision based on the stimuli. However, this differs substantially from typical real-life situations in which individuals actively seek out the information that they consider relevant to their choices. When making a natural choice, often one first determines whether and from which source to obtain information (e.g., to which stimulus to attend) and only then decides which action to take based on the information (Raiffa and Schlaifer 1961; S. C.-H. Yang, Wolpert, and Lengyel 2016; Gottlieb 2018; Braunlich and Love 2022). The sampling of decision-relevant (instrumental) information supports adaptive behaviors in humans and other animals (Gottlieb and Oudeyer 2018), and its disturbances are linked to psychopathology (Hauser et al. 2017; Baker et al. 2019), underscoring the importance of understanding its mechanisms.

In natural settings, information is gathered through active sensing behaviors—for example, when one attends to or looks at a visual cue (Tatler et al. 2011; S. C.-H. Yang, Wolpert, and Lengyel 2016)—or, alternatively, through explicit purchase decisions—for example, when a firm employs a consultant or a physician orders a medical test. Studies of information demand have primarily tested the latter scenario, using tasks in which participants are given a description of a situation and are asked to make a decision about whether or how much information to request in the situation (Furl and Averbeck 2011; Filimon et al. 2020; Kaanders et al. 2021; Gottlieb 2023). Neuroimaging investigations have focused on the value of information (VOI)— the extent to which obtaining information increases the rewards expected from future actions and choices (Raiffa and Schlaifer 1961; Howard 1966)—and showed that VOI is encoded in value and executive areas including the striatum, ventromedial prefrontal cortex, the dorsolateral prefrontal cortex, and the anterior cingulate cortex (Kobayashi and Hsu 2019; Kobayashi et al. 2021).

An open question, however, concerns the probabilistic computations that precede VOI estimation. In a decision-theoretic framework, VOI depends on both the rewards of a choice and, crucially, on the expected information gain associated with information gathering. Expected information gain, in turn, is the improvement in decision certainty that the decision-maker can expect to obtain by gathering information. A simple measure of expected information gain is *probability gain* (PG), defined as the difference between the decision maker’s prior certainty (PC)—the certainty about making the correct final choice before gathering information—and their *expected posterior certainty* (EPC)—the certainty that is *expected* after gathering information (Raiffa and Schlaifer 1961; Fischhoff and Beyth-Marom 1983; Baron 1985; Braunlich and Love 2022). PG describes humans’ demand for instrumental information in two-alternative inference tasks (Baron 1985; Nelson 2005; Nelson et al. 2010), but its neural mechanisms are not well understood.

A particularly challenging step in computing PG is the ability to prospectively reason about EPC—i.e., estimate the certainty that one expects to obtain after gathering information (Raiffa and Schlaifer 1961; Braunlich and Love 2022). To understand the neural correlates of EPC and related quantities, we used fMRI in a task in which participants placed bids revealing their willingness to pay for information relevant to a binary choice. On each trial, participants were informed about three quantities that were relevant to the normative bid: the prior probability that a decision alternative was correct, the diagnosticity of the information (the probability that it would correctly specify the correct alternative), and the penalty for an incorrect choice. By independently manipulating the three quantities, we distinguished between reward value and probabilistic computations of information gains. We show that the participants’ bids had independent sensitivity to PC and EPC, consistent with normative theories postulating that these quantities are combined to estimate PG. We also show that EPC was decoded with above-chance accuracy from the multivoxel activity in three regions in the right posterior parietal and extrastriate cortices. In these areas, the representation of EPC overlapped anatomically with representations of bids, PC, and PG that involved distinct activity patterns. The findings reveal candidate neural substrates for encoding expected information gain as a step that precedes but is distinct from the assignment of instrumental value to information.

## Results

### The Willingness to Pay Task

Participants (*N* = 23) underwent fMRI scanning while performing a task in which they bid for information relevant to an incentivized choice (**Figure 1A**). The task was a modified version of the “beads” task (Huq, Garety, and Hemsley 1988; Furl and Averbeck 2011; van der Leer et al. 2015; Baker et al. 2019), in which participants were told that each trial had a hidden state—a *portrait gallery* containing more pictures of faces than scenes or a *landscape* gallery containing more pictures of scenes than faces. Rather than asking participants to infer the gallery’s identity on each trial, as is customary, we first presented them with 120 trials in which they bid to receive additional information should they be asked to make the inference. Next, we randomly selected one trial from those the participants bid, delivered information with a probability that was proportional to the bid, asked participants to guess the gallery type, and delivered a payoff that depended on the accuracy of this guess (**Figure 1**).

**Figure 1.**
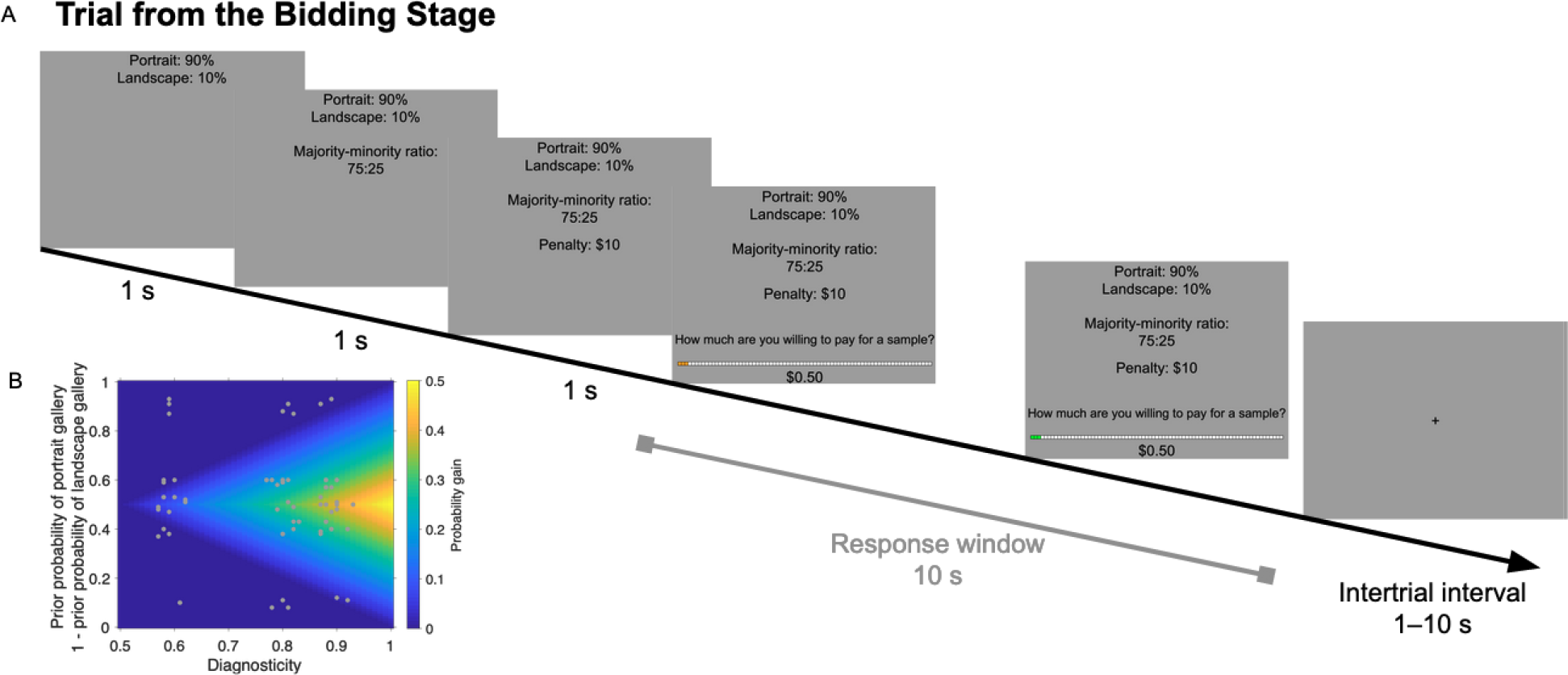
Task A: Structure of a bidding trial. On each trial, participants saw the complementary prior probabilities that the hidden gallery was a portrait or landscape gallery, the diagnosticity of a sample picture from the gallery (i.e., its evidence strength represented by the ratio of majority to minority pictures in the hidden gallery), and the amount the participant would be penalized from their endowment if the trial were realized for payment and they incorrectly guessed the hidden gallery. These quantities could appear in a variety of spatial or temporal orders. The participant then placed a bid for a sample picture from the hidden gallery, followed by a variable 1–10 s intertrial interval. **B: The distribution of prior probabilities and diagnosticities shown in the scan session** (white dots). The background color indicates PG, which increases with diagnosticity and decreases with PC (i.e., increases as the prior probabilities approach 0.5).

Our focus was on how the participants’ willingness to pay for information varied as a function of the context, as defined by three quantities that were relevant to the normative bid and were conveyed in words and numbers (**Figure 1A**). One quantity was the *penalty* for making an erroneous guess, a second quantity was the *prior probability* of a gallery type, and the third quantity was the *diagnosticity* of the information (the probability that the information, if given, would indicate the correct gallery type, conveyed as the ratio of pictures in the majority versus minority category; see **Methods**, Baker et al. (2019), and Furl and Averbeck (2011)). Importantly, the three quantities were statistically dissociated, allowing us to distinguish their influence on the bids. The penalty was randomly selected to be $10 or $20, while prior probability and diagnosticity were randomized independently to tile the probability space as shown in **Figure 1B**.

### Behavior

Our auction and payoff procedures incentivized participants to place a bid commensurate with the *value of information* (VOI), which was positively related to penalty and diagnosticity, and negatively related to the prior certainty (PC, the greater of the hidden gallery’s complementary prior probabilities; see **Methods, Equation 1**).

At both the group and individual levels, bids had a significant positive relationship with the normative VOI (**Figure S1**). A linear mixed-effects model (**Equation 14**) confirmed this result, yielding fixed-effects (group-level) coefficients that were negative for PC and positive for diagnosticity (**Figure 2A, Table 1**, the “Base Model”). The results were robust at the individual level, as the vast majority of participants had negative coefficients for PC and positive coefficients for diagnosticity and penalty (**Figure 2A**, gray dots; respectively, 22, 21, and 15 out of 23). These results, which were obtained for the scan session, were consistent with the same participants’ behavior during the prescan session and were replicated in two different cohorts who performed the task only outside of the scanner (**Figure 2A**, triangles, **Table 1**). The initial slider position was included as a nuisance regressor and produced negligible effects, ruling out sensorimotor artefacts (**Figure S2A**). Thus, participants understood the task and reliably placed bids that were consistent with normative VOI.

**Table 1.** Fixed-effects regression coefficients for the Base Model of participants’ bids (Equation 14) for every cohort (Main Cohort: *N* = 23, Cohort 1: *N* = 21, Cohort 2: *N* = 15). *DF*: Degree of freedom (from Satterthwaite approximation (Luke 2017)). *p*: *p*-value.

| Regressor | Cohort | Coefficient | Standard error | *T*-statistic | DF | *p* |
| --- | --- | --- | --- | --- | --- | --- |
| Prior certainty | Main, scan | −1.29 | 0.16 | −7.88 | 23.00 | < 0.001 |
|  | Main, prescan | −1.44 | 0.16 | −9.12 | 23.00 | < 0.001 |
|  | 2 | −1.82 | 0.14 | −14.40 | 21.00 | < 0.001 |
|  | 3 | −1.50 | 0.19 | −7.87 | 15.00 | < 0.001 |
| Diagnosticity | Main, scan | 0.93 | 0.19 | 4.88 | 23.00 | < 0.001 |
|  | Main, prescan | 0.94 | 0.18 | 5.34 | 23.00 | < 0.001 |
|  | 2 | 0.72 | 0.18 | 3.91 | 21.00 | < 0.001 |
|  | 3 | 1.17 | 0.23 | 5.20 | 14.99 | < 0.001 |
| Penalty | Main, scan | 0.19 | 0.11 | 1.72 | 23.01 | 0.098 |
|  | Main, prescan | 0.40 | 0.13 | 3.11 | 23.01 | 0.005 |
|  | 2 | 0.44 | 0.10 | 2.17 | 20.97 | < 0.001 |
|  | 3 | 0.37 | 0.09 | 4.08 | 15.00 | < 0.001 |

**Figure 2.**
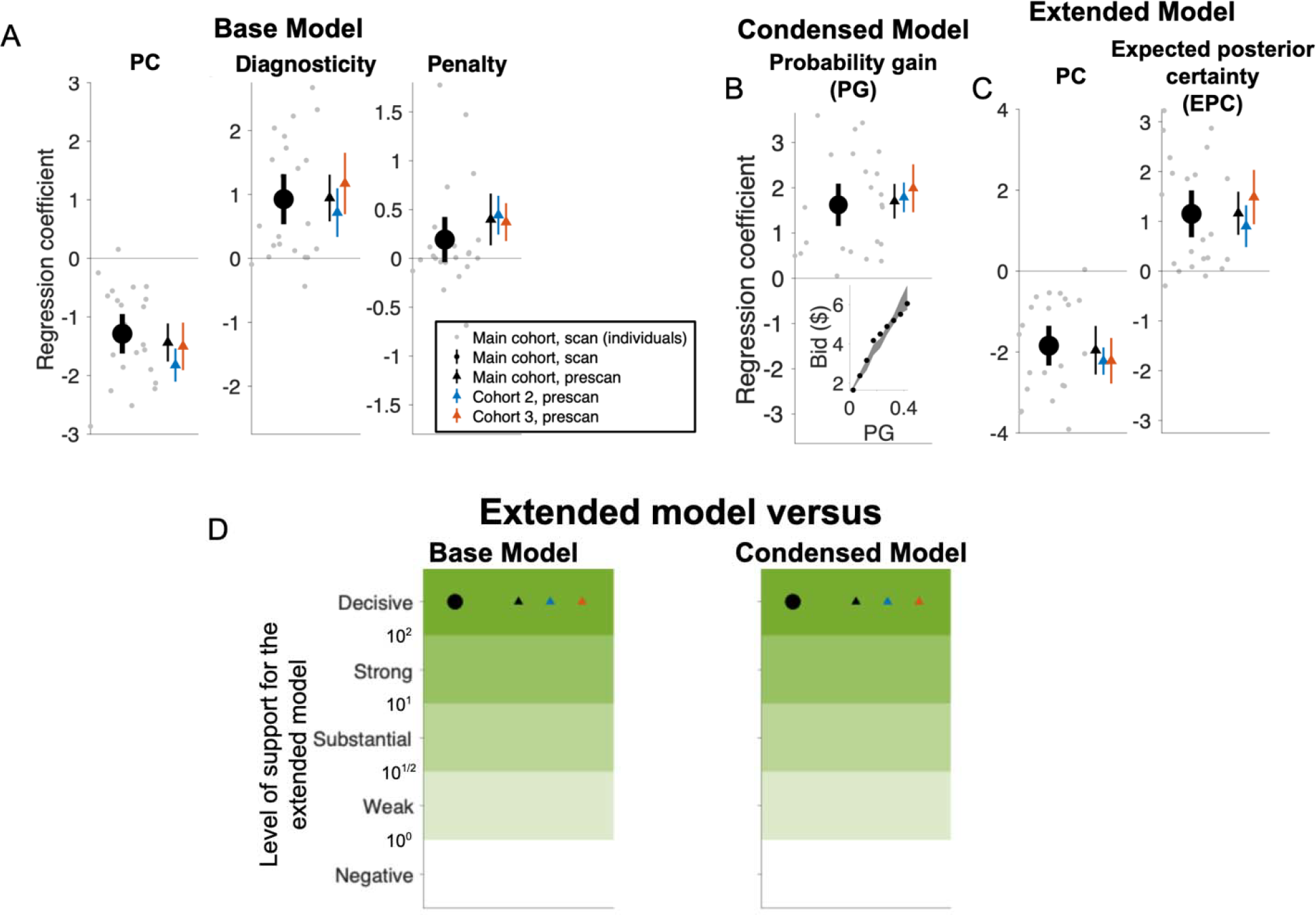
Participants’ bids are consistent with VOI. A–C: Coefficients from the three models of the bids. A: Base Model incorporating the variables shown on screen. Symbols show results from different participant groups. The large black dots show the fixed-effects (group-level) coefficients from the main cohort during the scan session (*N* = 23). The small gray dots show the random (individual) effects for the same group. The black triangles show the fixed effects from the main cohort during the prescan session (*N* = 23), and the blue and gray triangles show two additional cohorts that were tested only behaviorally (*N* = 21 and *N* = 15, respectively). Error bars represent 95% confidence intervals. Detailed statistics on the fixed-effects coefficients are in **Table 1. B: The Condensed Model based on PG**. Same format as in **A**. *Inset:* Visualization of the increase in bids with PG. Each point is the mean submitted bid across all completed trials, binned by PG. The shaded area shows the Condensed Model’s predictions of submitted bids for each bin (mean and 95% confidence intervals). Detailed statistics on the fixed-effects coefficients are in **Table 2. C: The Extended Model based on PC and EPC**. Same format as in **A** and **B**. Detailed statistics on the fixed-effects coefficients are in **Table 3. D: Model comparisons using Bayes factors**. The symbols show the Bayes factors (BF) comparing the extended model to the base model (left) and the condensed model (right) for the cohorts in **A–C**. For ease of presentation, BF values were divided into categories showing negative, weak, substantial, strong, and decisive support for the extended model (respectively, BF < 10^0^, 10^0^ ≤ BF < 10^1/2^, 10^1/2^ ≤ BF < 10^1^, 10^1^ ≤ BF ≤ 10^2^ and 10^2^ < BF (Kass and Raftery 1995). All cohorts showed decisive evidence in favor of the Extended Model over both the Base Model (Main Cohort scan session: 1.30×10^76^, Main Cohort prescan session: 1.15×10^31^, Cohort 2: 1.13×10^3^, Cohort 3: 8.36×10^66^) and the condensed model (Main Cohort scan session: 1.69×10^109^, Main Cohort prescan session: 5.93×10^99^, Cohort 2: 4.34×10^168^, Cohort 3: 1.25×10^53^).

While these results show that participants are sensitive to the information shown on the screen, VOI depends on several quantities that are derived from this information. Specifically, VOI scales with probability gain (PG), which is the difference between PC and expected posterior certainty (EPC) and, in turn, EPC is derived from prior probability and diagnosticity (**Equation 4**). To examine if participants estimated these quantities, we fit their bids to two additional models: the *Condensed Model* accounting for the effect of PG (**Equation 15**) and the *Extended Model* separately capturing the effects of PC and EPC (**Equation 16**). Consistent with the Base Model, the Condensed Model produced positive fixed effects for PG (**Figure 2B**), and the Extended Model yielded positive fixed-effects coefficients for EPC and negative fixed-effects coefficients for PC (**Figure 2C**). All fixed-effects coefficients were highly significant and robust across groups (see detailed statistics in **Table 2** and **Table 3)**, were highly consistent at the individual level (gray dots in **Figure 2B–C**) and could not be explained by sensorimotor artifacts (see **Figure S2** for all the coefficients for all three models).

**Table 2.** Fixed-effects regression coefficient for the Condensed Model of participants’ bids (Equation 15) for every cohort (Main Cohort: *N* = 23, Cohort 1: *N* = 21, Cohort 2: *N* = 15). *DF*: Degree of freedom (from Satterthwaite approximation (Luke 2017)). *p*: *p*-value.

| Regressor | Cohort | Coefficient | Standard error | *T*-statistic | DF | *p* |
| --- | --- | --- | --- | --- | --- | --- |
| Probability gain | Main, scan | 1.62 | 0.23 | 7.18 | 23.00 | < 0.001 |
|  | Main, prescan | 1.70 | 0.19 | 9.19 | 22.99 | < 0.001 |
|  | 2 | 1.79 | 0.16 | 11.33 | 21.00 | < 0.001 |
|  | 3 | 1.99 | 0.25 | 8.01 | 15.00 | < 0.001 |

**Table 3.** Fixed-effects regression coefficients for the Extended Model of participants’ bids (Equation 16) for every cohort (Main Cohort: *N* = 23, Cohort 1: *N* = 21, Cohort 2: *N* = 15). *DF*: Degree of freedom (from Satterthwaite approximation (Luke 2017)). *p*: *p*-value.

| Regressor | Cohort | Coefficient | Standard error | *T*-statistic | DF | *p* |
| --- | --- | --- | --- | --- | --- | --- |
| Prior certainty | Main, scan | -1.84 | 0.24 | -7.75 | 23.00 | < 0.001 |
|  | Main, prescan | -1.95 | 0.20 | -9.90 | 22.99 | < 0.001 |
|  | 2 | -2.22 | 0.16 | -13.71 | 21.00 | < 0.001 |
|  | 3 | -2.22 | 0.26 | -8.38 | 15.00 | < 0.001 |
| Expected posterior certainty | Main, scan | 1.15 | 0.23 | 5.05 | 23.00 | < 0.001 |
|  | Main, prescan | 1.16 | 0.21 | 5.55 | 23.00 | < 0.001 |
|  | 2 | 0.90 | 0.20 | 4.46 | 21.00 | < 0.001 |
|  | 3 | 1.49 | 0.26 | 5.80 | 15.00 | < 0.001 |

Importantly, model comparisons decisively favored the extended model over both the condensed and base models (**Figure 2D)** with Bayes factors far exceeding 10^3^ in all the cohorts (see legend for specific values). Moreover, in the extended model, the average difference between the absolute value of each participant’s coefficients for PC and EPC was significantly positive (scan session: 2.52, *T*(22) = 2.60, SE = 0.97, *p* = 0.016, *N* = 23, paired *T*-test), suggesting that participants integrated PC and EPC with unequal weights that relatively underweighted EPC. Thus, rather than merely combining the quantities presented on the screen, participants estimated the posterior certainty they expected to have after gathering information and weighted it separately from their initial uncertainty when they bid for information. This motivates the investigation of a neural representation of EPC as a neural basis of prospective Bayesian inference.

### Neural Representations of Probabilistic Variables Relevant for VOI

To identify neural representations of the variables involved in calculating the bids, we applied multi-voxel pattern analysis (MVPA) to blood-oxygen-level-dependent (BOLD) signals during the response window of each trial in the bidding phase (**Figure 1A**). We used support vector regression and leave-one-run-out (four-fold) cross-validation and measured decoding accuracy for bid, PG, PC, and EPC as the Fisher *z*-transformed correlation between the predicted and actual values of the variable (Fisher 1921; 1915; Görgen and Hebart 2022).

Considering that EPC is a numerical representation of prospective Bayesian inference, we first used a whole-brain searchlight analysis to identify clusters that showed significantly above-chance decoding of EPC with no cross-decoding of slider displacement or of bid, PG, and PC. This identified three clusters that met these criteria located, respectively, in the right occipital fusiform gyrus, right occipital pole, and right intraparietal sulcus (IPS)/extrastriate cortex (**Figure 3A–B**, and **Table S1**). Decoders trained to read out EPC in these clusters produced no significant cross-decoding of slider displacement, providing no credible evidence that they encoded visual or motor events (**Figure 3C**). Moreover, the decoders produced no significant cross-decoding of bids (**Figure 3D**, left) despite the fact that EPC was correlated with bids as a necessary corollary of good task performance (**Figure 2C**), suggesting that they represented EPC independently of information value. The clusters also did not cross-decode PG (**Figure 3D**, middle), despite the fact that PG is the difference between EPC and PC (**Equation 5**). Finally, the clusters also did not decode PC (**Figure 3D**, right), which, after excluding a small subset of high-leverage trials that were high in both variables, was uncorrelated with EPC. Thus, the three cortical clusters conveyed information about EPC independent of slider position, PG, PC, or the value of information as reflected in bids.

**Figure 3.**
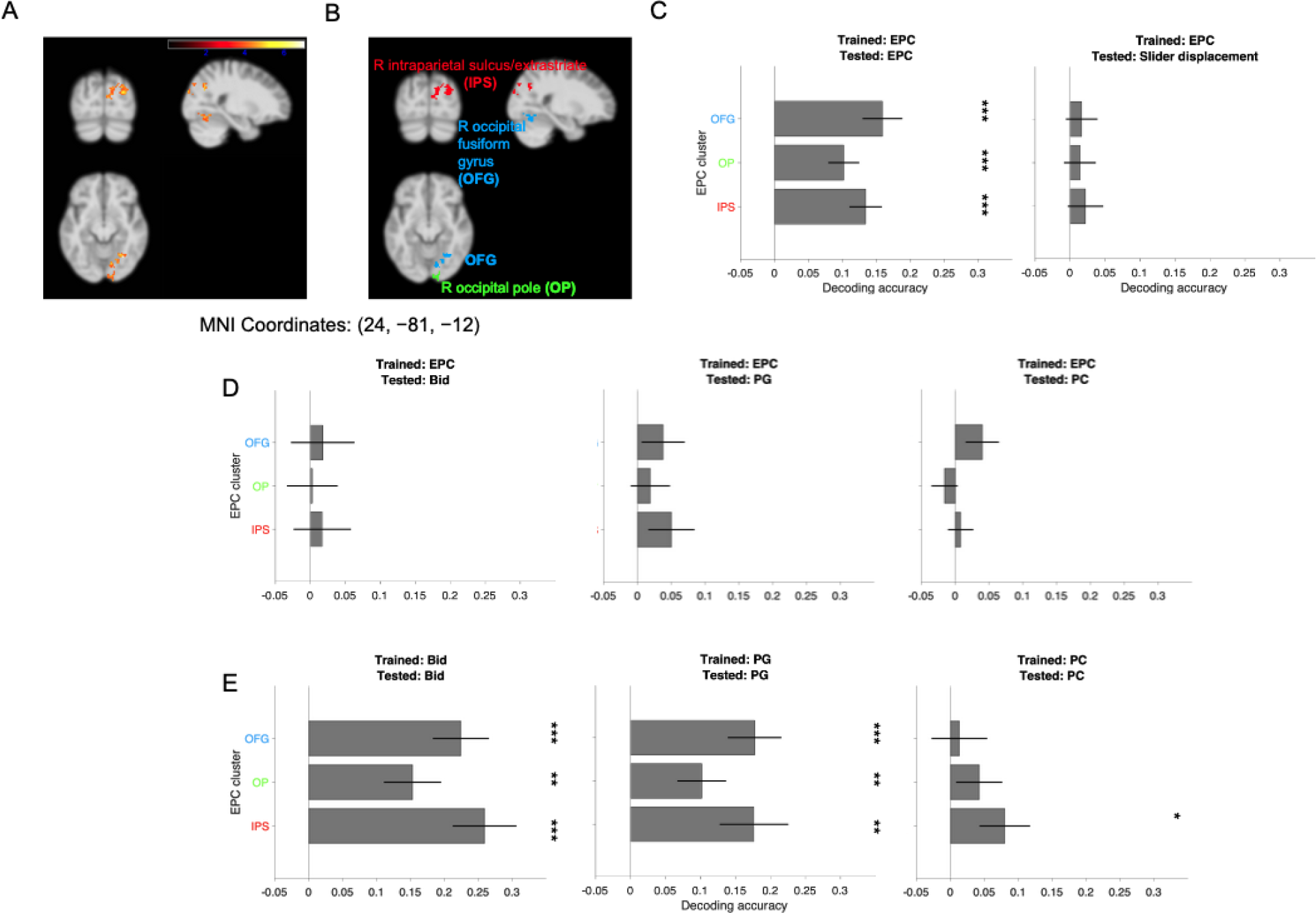
Distinct multivoxel representations of expected posterior certainty (EPC), probability gain (PG), prior certainty (PC), and bid in the posterior parietal and extrastriate cortices. **A:** Thresholded *T*-statistic map of significant clusters of activation in which EPC could be decoded above chance, as identified by a whole-brain searchlight analysis (cluster-defining height threshold: *p* < 0.001; cluster-level familywise error rate correction threshold: *p* < 0.05). The anatomical template was smoothed at FWHM = 5×5×5 mm for visualization purposes (Poldrack, Mumford, and Nichols 2011, 173). **B:** The labeled significant clusters in which EPC could be decoded: right occipital fusiform gyrus (OFG)/cerebellum, light blue; right occipital pole, green; and right intraparietal sulcus (IPS) and adjacent extrastriate cortex, red. **C–E: Decoding results from the 3 clusters**. Each panel shows the decoding accuracy of ROI-wise decoders that were trained and tested on the quantities noted in the panel title. Each bar shows the mean decoding accuracy and standard errors across 23 participants. ***: *p* < 0.001, **: 0.001 ≤ *p* < 0.01, *: 0.01 ≤ *p* < 0.05.

We next asked whether these clusters may have had representations of bid, PG, and PC that anatomically overlapped with the representation of EPC but involved distinct activity patterns. Indeed, when we trained new decoders to decode bid and PG, we found significant above-chance decoding in all 3 ROIs (**Figure 3D**, left, middle) and, when we trained new decoders to decode PC, we found significant above-chance decoding in the right IPS/extrastriate cluster (**Figure 3D**, right). The lack of cross-decoding documented in **Figure 3C** makes it unlikely that these results merely reflected correlations between these variables and EPC. Therefore, the clusters encoding EPC multiplex distinct neural representations of bid, PC, and PG.

Separate whole-brain searchlights identified additional clusters that encoded PC and PG (**Figure S3**). These clusters showed no cross-decoding of slider position, showing that they were unlikely to reflect sensorimotor confounds, nor significant cross-decoding of EPC (**Figure S3**). However, the clusters did show significant cross-decoding between PG and PC and between each variable and the participants’ bids, making it difficult to pinpoint precisely which variable they encoded toward computing the bid.

## Discussion

We used behavioral testing and fMRI to investigate the mechanisms by which people estimate expected information gain when assigning value to information. We provide evidence that, consistent with value of information (VOI) theory, participants’ estimates of VOI were informed by the difference between expected posterior certainty (EPC) and prior certainty (PC). Moreover, we show that portions of the right posterior parietal and extrastriate cortex conveyed distinct multi-voxel representations of EPC and PC, which spatially overlapped with each other, as well as with distinct representations of PG and the participants’ bids. The results support the hypothesis that participants prospect about future posterior probabilities and information gains when estimating the instrumental value of information and reveal neural substrates underlying this process.

An important feature of our task is that participants did not learn through repeated experience but made one-shot decisions about the value of information based on quantities explicitly shown on the screen: prior probability, diagnosticity, and penalty. While this differs from some studies of Bayesian inference that allow learning based on feedback (Soltani and Wang 2010; Kira, Yang, and Shadlen 2015; Ting et al. 2015; Soltani et al. 2016), it closely follows studies of information gathering that have typically relied on one-shot decisions based on a description of behavioral context (Kobayashi and Hsu 2019; Filimon et al. 2020; Kobayashi et al. 2021; Gottlieb 2023). A second important task feature is that we limited participants to a single additional sample rather than allowing them to request multiple samples. While this distinguishes our approach from studies examining how people terminate sampling (i.e., decide *how much* information to gather before making a choice (Edwards 1965; Huq, Garety, and Hemsley 1988; Roitman and Shadlen 2002; Furl and Averbeck 2011; Hanks, Kiani, and Shadlen 2014; Baker et al. 2019; Kaanders et al. 2021; Ashinoff et al. 2022)), it allowed us to understand with greater experimental control how participants prospect about information gains over a single time step (e.g., avoiding systematic distortions and noise that may gradually accumulate over samples (Ashinoff et al. 2022)).

Our results support the idea that participants prospect about future certainty as noted above, and also show that, rather than directly comparing PC to EPC with equal weights, they afforded greater weight to PC relative to EPC. A possible explanation for this differential weighting is that EPC is derived through more complex computations making it more vulnerable to probability underweighting, a known phenomenon during judgments from described probabilities (Gonzalez and Wu 1999; Trepel, Fox, and Poldrack 2005; Garcia, Cerrotti, and Palminteri 2021). Alternatively, participants may underuse EPC because perhaps prospection itself is costly. These factors, in turn, may explain why participants prospect over a limited time horizon when allowed to take sequential samples (Braunlich and Love 2022), as can be examined in future research.

Our approach also generated new insights into the neural substrates underlying information gathering. The encoding of probabilistic variables that we found in the posterior parietal cortex is consistent with multiple studies that have implicated this area in probabilistic reasoning. In tasks in which participants make Bayesian inferences based on given (exogenous) information, the human posterior parietal cortex tracks prior probability (Mulder et al. 2012), likelihood (d’Acremont, Fornari, and Bossaerts 2013; d’Acremont, Schultz, and Bossaerts 2013), likelihood uncertainty (Ting et al. 2015), and posterior probability (Singletary, Gottlieb, and Horga 2021), while monkey parietal neurons encode posterior probability or expected rewards (Huk and Shadlen 2005; Kira, Yang, and Shadlen 2015; T. Yang and Shadlen 2007). In tasks in which participants endogenously select information, monkey parietal neurons encode diagnosticity (Foley et al. 2017) and prior uncertainty (Horan, Daddaoua, and Gottlieb 2019; Li et al. 2022) and the human parietal cortex tracks the propensity to sample information relevant for learning a category boundary (Furl and Averbeck 2011).

Our findings extend these reports by showing that the human parietal cortex and extrastriate areas multiplex information about probabilistic variables—of EPC, PC, and PG—that are distinct from variables representing information value. Thus, our results support the idea that information gathering has separate probabilistic and value-based components, as proposed based on both behavioral (Braunlich and Love 2022) and neural (Silvetti et al. 2023) results. This result is consistent with studies showing that monkey parietal neurons carry dissociable signals of prior uncertainty and rewards (Horan, Daddaoua, and Gottlieb 2019; Li et al. 2022). The findings are also consistent with previous studies proposing that VOI is encoded in areas that are distinct from the EPC clusters we identified here, and include the human striatum, dorsolateral prefrontal cortex, and ventromedial prefrontal cortex (Kobayashi and Hsu 2019; Kobayashi et al. 2021). Moreover, our findings that additional clusters non-specifically encode PC, PG, and bids, suggesting the possibility, which can be tested in future research, that probabilistic and value quantities are integrated into a single code for driving the bids. Thus, our results bring granular insights into the distinct neural mechanisms of probability and reward estimation during active information gathering.

## Methods

### Participants

Forty-four healthy, right-handed participants (17 female) were recruited through fliers posted on the Columbia University campus and through the recruitment system for the Columbia Business School Behavioral Research Lab. This pool consisted of Columbia University students, other Columbia affiliates, and affiliates of other universities in the New York Metropolitan Area, and they did not report any psychiatric or neurological disorders. Participants first completed a session outside of the scanner (prescan session); 14 participants were not allowed to advance to the scan session because their responses during the prescan session reflected disengagement or lack of comprehension (see “Performance-Based Exclusion Criteria”). Another participant was excluded because of excessive motion inside the MRI scanner, and 6 participants met the advancement criteria but withdrew from the scan session. As a result, the Main Cohort consisted of 23 participants (8 female).

We also recruited 19 participants (13 female) through the same methods to complete the experiment outside of the scanner. Fourteen participants (10 female) met the comprehension criteria and were included in the Cohort 2. Added to Cohort 2 were the participant who was excluded because of excessive motion and the 6 participants who withdrew from scanning in the Main Cohort.

Before developing the main experimental session, we recruited 23 participants (9 female) through the same methods for a pilot session to be completed outside of the scanner. Fifteen of these participants (7 female) met the comprehension criteria and were included in Cohort 3. Experimental procedures were approved by the Columbia University Institutional Review Board, and all participants provided signed informed consent.

### Experimental Sessions

#### Prescan Session

The prescan session was administered on a computer outside of the scanner. Participants viewed a narrated slideshow on the instructions for the Willingness to Pay (WTP) Task, the main task of the experiment. They were also administered comprehension quizzes on the instructions, which they had to pass before proceeding (**Performance-Based Exclusion Criteria**). After passing the instructions quiz, participants completed 10 practice trials of the WTP Task to familiarize themselves with the relationship between their bids, the receipt of a sample picture, decision accuracy, and ultimately, their earnings, all while avoiding overtraining. Each practice trial was followed by a corresponding mock payout trial to show participants what they could have earned from that trial in the main task based on their submitted bid and their guess of the identity of the hidden gallery if the trial had been chosen for payout; however, these practice trials did not affect the participants’ earnings. Then, participants completed the WTP Task. Lastly, their performance was evaluated to determine if they met the remaining performance criteria to advance to the scan session; if not, they were removed from the study.

#### Scan Session

Participants watched a summarized version of the instructions slideshow before completing the WTP Task in the MRI scanner.

#### The Willingness to Pay Task

The Willingness to Pay (WTP) Task was a modified “bookbag-and-poker-chip” (Peterson and Miller 1965; Phillips and Edwards 1966; Bar-Hillel 1980; Gigerenzer, Hell, and Blank 1988; Benjamin 2019) (or “beads” (Huq, Garety, and Hemsley 1988; Furl and Averbeck 2011; van der Leer et al. 2015; Baker et al. 2019; Kobayashi et al. 2021)) task developed to measure people’s demand for instrumental information. On the task, people needed to correctly infer the identity of a hidden state depicted as a museum gallery to avoid a penalty. The task consisted of a *Bidding Stage* followed by a *Payout Stage*. At the beginning of each session, the participant was given a $30 endowment. Then, during the Bidding Stage, the participant placed bids for a sample picture from the hidden museum gallery that could increase the accuracy of their inference. During the Payout Stage, one bidding trial was drawn at random to be realized to determine the participant’s payout. The bid on the realized trial was applied to a computer-automated auction that ensured that the probability of receiving a sample increased with the bid such that a higher bid corresponded to higher demand for the sample (**The auction and the expected value– maximizing bid**). The bid would be withdrawn from the endowment if the participant won the auction on the realized trial, and a penalty would be withdrawn if the participant’s inference was incorrect.

#### Bidding Stage

The Bidding Stage consisted of 120 trials divided evenly into four runs. On each trial, participants bid for one sample picture from the hidden gallery that could help them better infer whether it was a *portrait gallery* that contained more pictures of faces than scenes or a *landscape* gallery that contained more pictures of scenes than faces. Before bidding, participants viewed the *prior probability* that the hidden gallery was a portrait or a landscape gallery; the *diagnosticity*, or predictive validity, of a sample picture; and the *penalty* that they would lose if the trial were realized for payout and they incorrectly guessed the gallery.

To prevent behavioral artifacts from serial trial effects, we truthfully told participants that each bidding trial was independent from all other bidding trials, and the identity of a bidding trial’s hidden gallery was never revealed during the Bidding Stage.

*Trial display*. The prior probability of the hidden gallery was displayed as a percentage chance for the portrait and landscape options. The diagnosticity was displayed as the majority-to-minority ratio of picture types in the hidden gallery (e.g., 60:40). Participants were also shown the *penalty* that they could lose from the endowment if the trial were chosen for payout (**Payout trial**).

A trial began with the prior probability, diagnosticity, or penalty appearing (trial components) over a gray background (**Figure 1A**). The prior probability, diagnosticity, and penalty appeared one at a time with the first component appearing at the instant of trial start and the succeeding components following the previous component by 1 s (**Figure 1A**). The trial components’ spatial order of appearance was stable throughout the prescan and scan sessions but counterbalanced by participant so that participants could expect the information to be in the same place while allowing us to control for potential effects of spatial order. The trial components’ temporal order of appearance was randomized by trial to control for potential primacy and recency effects.

*Response*. Participants completed a trial by reporting their bid for a sample picture from the hidden gallery by using a trackball to move a slider that appeared at the bottom of the screen 1 s after the last trial component. The initial slider position was randomized on each trial to reduce the correlation between slider movement and the bid—facilitating the separation of the potentially confounding effect of slider movement from the task variables of interest—and to discourage participants from anchoring to any one reported bid. (Randomizing the initial slider position reduces the correlation between slider displacement and bid from nearly 1 to 0.58 across all completed trials in the scan session.) The slider remained on screen for 10 s (“response window,” **Figure 1A**). We chose a response window of 10 s because it was the shortest response window that captured approximately 80 percent of responses from 80 percent of participants during piloting. The selected bid was indicated by the amount of the slider from left to right that was highlighted in orange and by an explicit amount in dollars below the slider. Both these indicators were updated in real time. The slider was divided into 77 discrete bins, increasing in steps of $0.13 from $0 on the left to $9.88 on the right. We chose these increments to discourage participants from anchoring to “round” numbers (e.g., multiples of $1 or $2.50). The participant confirmed their response by clicking a button on the trackball, after which the highlighted section of the slider would change colors from orange to green to indicate that the response had been recorded. The screen remained unchanged until the end of the response window plus 0.5 s. If the participant did not submit a posterior probability estimate within the 10-s response window, instead, the slider would freeze for 0.5 s and the percentage below the slider would be replaced by text reading, “Bid not submitted.” To encourage participants to respond within the response window, participants were truthfully warned that if a response were missing from a trial that happened to be chosen for payout, they would automatically lose that trial’s penalty. Across all participants in the Main Cohort during the scan session, only 23 of the 2,760 presented trials (0.8%) had omitted responses, with 7 participants missing one trial, 4 participants missing two trials, 1 participant missing three trials, and 1 participant missing five trials.

*Intertrial interval*. Each bidding trial was followed by an intertrial interval during which a small, black fixation cross appeared over the gray background (**Figure 1A**). To maximize the efficiency of parameter estimation for the general linear models in the fMRI analysis, the duration of each intertrial interval was drawn from an exponential distribution with mean 3.5 s, truncated with a lower bound of 1 s and an upper bound of 10 s (Hagberg et al. 2001).

*Selection of parameters for bidding trials*. To determine the set of prior probabilities and majority-minority ratios used for the bidding trials in each session, we randomly sampled 60 trials from discrete bins that we established for prior probability (0.1, 0.4, 0.5, 0.6, and 0.9, arbitrarily chosen as the prior of the portrait gallery) and majority–minority ratio (60:40, 80:20, and 90:10). Majority–minority ratios represented diagnosticity θ, which was defined on the interval 0.5 ≤ θ ≤ 1 and corresponded to the numerator of the majority–minority ratio divided by 100. A random jitter (−0.03, −0.02, −0.01, 0, 0.01, 0.02, or 0.03) was then added to each prior probability and diagnosticity with equal probability. A “true” hidden gallery was assigned to each trial based on the prior probability of the portrait gallery (e.g., if the prior probability was 0.6, there was a 60% chance the trial’s hidden gallery would be a portrait gallery and a 40% chance it would be a landscape gallery). If the trial were realized for payout, its sample picture was assigned to signal the hidden gallery with a probability equal to the trial’s diagnosticity (e.g., on a trial on which the hidden gallery was a portrait gallery and the diagnosticity was 0.6, there was a 60% chance that the sample picture would be a face). These 60 trials were duplicated for each condition of error penalty ($10 or $20). The order of the trials was then randomly permuted, and the session was separated into four runs of 30 trials each. **Figure 1B** displays the prior-diagnosticity combinations for the scan session. Every participant within a cohort completed the same session(s).

#### Payout trial

After the Bidding Stage was complete, one bidding trial was chosen at random with equal probability to be realized to determine the participant’s payment. This trial was displayed along with its submitted bid from the Bidding Stage. If the participant had failed to submit a bid on that trial, the participant was notified that the error penalty would be automatically subtracted from their endowment, and the session would end.

Otherwise, the bid was submitted to an auction for the sample picture. If the participant had bid enough to win the auction, the bid would be subtracted from their endowment, and they would receive one sample picture, randomly drawn from the hidden gallery, to help them decide the hidden gallery’s identity (in addition to the prior probability of the hidden gallery and the diagnosticity of a sample picture). If the participant had not bid enough to win the auction, the bid would not be subtracted from their endowment, but they would have to decide using only the prior probability of the hidden gallery.

After deciding the identity of the hidden gallery, the penalty would be subtracted from the participant’s endowment if and only if their choice was incorrect. Hence, the payment for the WTP Museum Task was the endowment minus the bid (if and only if they received a sample) and minus the penalty (if and only if they chose the incorrect gallery).

#### The auction and the expected value–maximizing bid

We elicited participants’ valuation of the sample through a first-price auction. On the realized trial, the computer chose but did not reveal a secret price for the sample picture from a random uniform distribution between the minimum and maximum possible bids ($0 and $9.88, respectively). If the participant’s bid on the realized trial was greater than or equal to the secret price, the participant would receive the sample, but their bid would be subtracted from their endowment. If the participant’s bid was less than the secret price, the participant would not receive the sample, but nothing would be subtracted from their endowment. This procedure is inspired by the Becker-DeGroot-Marschak auction (Marschak, DeGroot, and Becker 1964) and is similar to a traditional auction for an item in which the highest bidder receives the item in exchange for their stated price. Hence, the participant must assess the risk of losing the penalty if they incorrectly guessed the hidden gallery against the potential cost of a sample picture that could decrease the chance of an incorrect guess.

This cost-benefit analysis for the WTP Task bids is mathematically tractable for an expected value maximizer. Intuitively, the expected value–maximizing bid in this auction increases with the expected information gain (EIG), or informativeness, of the sample—the greater the sample’s EIG, the more an agent should be willing to pay to receive a sample. To express the expected value–maximizing bid in terms of EIG, we measured EIG using *probability gain*, the extent to which the sample picture would increase the probability of correctly guessing the hidden gallery (Baron 1985; Nelson 2005). We chose probability gain because it has been shown to modulate people’s demand for instrumental information (Nelson et al. 2010) and because bids from the auction can be expressed as a simple linear function of it.

To estimate probability gain, one needs to compare the probability of correctly guessing the hidden gallery without the sample picture to the expected probability of correctly guessing the hidden gallery with the sample picture. We call the first component of probability gain the *prior certainty* of the hidden gallery (Pr(*C*)), which is the maximum of the hidden gallery’s prior probability distribution (**Equation 1**). Here, Pr(*F*) is the prior probability of the portrait gallery, and Pr(*S*) is the prior probability of the landscape gallery.

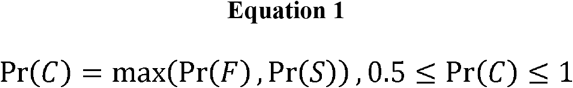

We call the second component of probability gain the *expected posterior certainty* (EPC) of the hidden gallery. Calculating the expected posterior certainty requires prospecting the future posterior probability distributions for a gallery type (*F* for the portrait gallery, *S* for the landscape gallery), taking the maximum of these posterior distributions conditional on receiving each type of sample picture (for a face picture, for a scene picture), and weighting each maximum by the marginal probability of the respective picture type.

The posterior probability of gallery *H* conditional on sample picture *D* is given by Bayes’ theorem in terms of the prior probability of the gallery (Pr(*H*)), the likelihood of receiving sample picture *D* conditional on the gallery (Pr(*H*)), and the marginal probability of the sample (Pr(*D*)) (**Equation 2**, where *H* stands for “hypothesis” and *D* for “data,” by convention). The prior probability of each gallery type is explicitly shown on a trial, while the likelihood and marginal probability of a sample type can be calculated from variables that are shown on a trial. The prospected likelihood is the diagnosticity (*θ*) if the sample picture is prospected to signal gallery *H* (i.e., when gallery *H* is the portrait gallery and the sample is prospected to be a face, or when gallery *H* is the landscape gallery and the sample is prospected to be a scene), while the likelihood is the complement of the diagnosticity (1 − *θ*) if the sample picture is prospected to signal the opposite gallery. The prospected marginal probability of sample *D* is the product of the diagnosticity of the sample and the prior probability of the gallery signaled by the sample plus the product of the complement of the diagnosticity and the complement of the prior probability signaled by the sample (**Equation 3**).

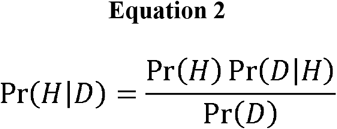

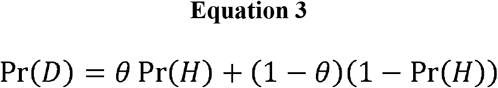

Therefore, we can use the marginal probability of each sample type and the posterior probability distribution for each gallery to calculate the expected posterior certainty of the hidden gallery (**Equation 4**).

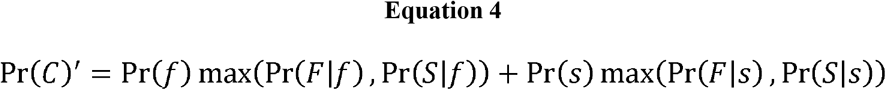

This is equivalent to the maximum of the prior certainty and the diagnosticity (Pr(*C*)’ = max(Pr(*C*), *θ*)).

Therefore, in terms of prospecting the expected certainty after receiving a sample picture, probability gain *G* is the EPC minus the prior certainty (**Equation 5**).

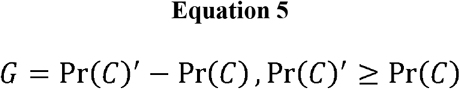

Probability gain can be equivalently expressed as a rectified function of the sample’s diagnosticity and the prior certainty of the hidden gallery (**Equation 6**).

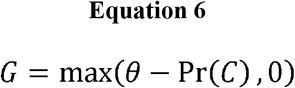

To calculate the bid *B*_𝔼max_ in this auction that maximizes the expected value of a trial, we first need to calculate the expected value of a trial 𝔼 in terms of the endowment, the penalty, Pr(*C*), and Pr(*C*)’. Expected value is the value of the outcome minus its cost. Since the value and cost of a trial depend on whether the agent receives a sample, which is a random event, let the value be 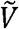, a random variable representing the value of the *outcome*, and let the cost of the outcome be *C*, a random variable representing the price the agent ultimately pays for the sample (**Equation 7**).

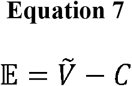

To calculate *E* over all the possible realizations of a trial, let us calculate the probability density function for receiving a sample. As stated earlier, the computer chooses a random price *X* in dollars from a uniform probability distribution on the interval of possible bids 0 ≤ *X* ≤ 9.88. Thus, the probability density function for the random variable is^1^

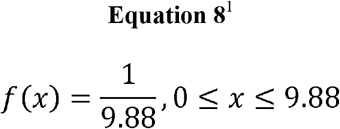

Now, let us take 𝔼 over all possible realiza *X:*

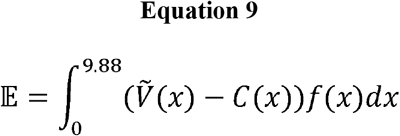

**Equation 9** can be decomposed into the utility of winning the auction (the left addend in **Equation 10**) and the utility of losing the auction (the right addend in **Equation 10**):

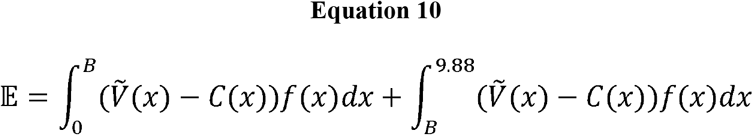

Now let us replace the random variables 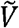 and *C* with the exact outcomes that they represent. To do so, first let us construct the tree of possible outcomes for a realized trial: at the first step, the agent may win or lose the auction for the sample picture, and at the second step, the agent may correctly or incorrectly guess the identity of the hidden gallery:

- When the agent wins the auction
  - When the agent correctly guesses the hidden gallery
    - 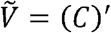 (because they win the full endowment, which is $30)
    - C=Pr(*C*)’*B*
  - When the agent incorrectly guesses the hidden gallery
    - 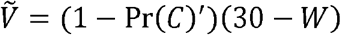
    - *C=*(*1-*Pr(*C*)’)*B*
- When the agent loses the auction
  - When the agent correctly guesses the hidden gallery
    - 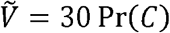
    - *C* = 0
  - When the agent incorrectly guesses the hidden gallery
    - 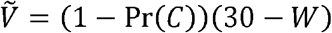
    - *C* = 0

Replacing 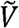, *C*, and *f* (*x*) in **Equation 10** with the above terms yields the expected value of a trial in terms of the endowment, the penalty, Pr(*C*), and Pr(*C*)*’* (**Equation 11**).

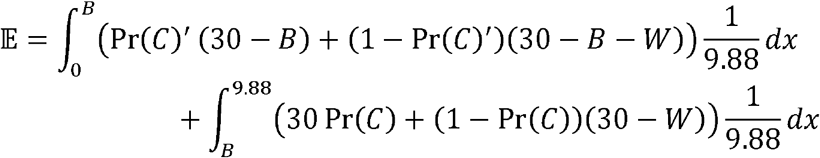

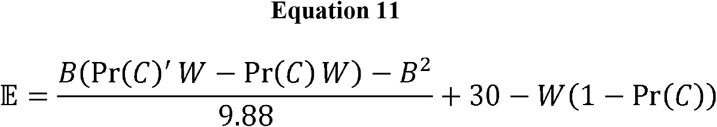

To find the expected value–maximizin (**Equation 12**), let us differentiate *E* with respect to the bid, set the derivate equal to 0, and solve for the bid *B* _*𝔼max*_:

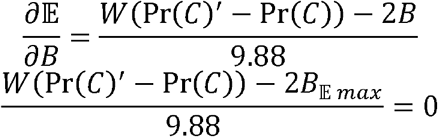

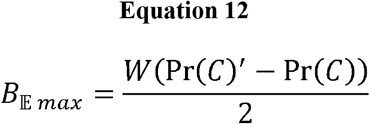

Note that the endowment does not affect the expected value–maximizing bid.

Recall that probability gain is the difference between EPC and prior certainty. Therefore, we can rewrite **Equation 12** in terms of probability gain *G* (**Equation 13**). We use both this form and the form in **Equation 12** to model participants’ bids in terms of expected information gain.

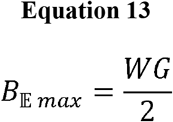

Since the WTP Task only accepts bids in bins (**Figure 1A**), on the real task, the expected value is maximized by submitting a bid as close as possible to the expected value–maximizing bid.

#### Performance-Based Exclusion Criteria

To ensure participant comprehension and engagement during the scan session, we assessed participants’ performance during the prescan session before we allowed them to advance to the scan session. Participants had to meet the following criteria pertaining to the WTP Task to advance to the scan session:

1. Task comprehension: Participants had to correctly answer at least 80 percent of the questions on a comprehension quiz on the task instructions.
2. Task completion: Participants could miss no more than 5 percent of the trials.
3. Minimal dependence of bids on expected information gain (EIG): Bids must have been significantly higher (α = 0.05, two-sample t-test assuming unknown and unequal variances) on trials with high probability gain (≥ 0.33) than on trials with low probability gain (≤ 0.1).

To measure participants’ intrinsic demand for instrumental information without extensive training, the criteria were designed to be lenient enough to respect variation in their pre-task strategies while excluding participants who disengaged from the task or who adopted strategies clearly consistent with misunderstanding the task.

#### Image Sets

Images of faces were selected from the CNBC Faces database by Michael J. Tarr, Center for the Neural Basis of Cognition and Department of Psychology, Carnegie Mellon University, http://www.tarrlab.org, funded by NSF award 0339122, used in Righi et al. (2012), and are available under a Creative Commons Attribution-NonCommercial-ShareAlike 3.0 Unported License. Images of scenes were selected from the database for Konkle et al. (2010)), available from the Computational Perception and Cognition Lab at MIT (http://olivalab.mit.edu/MM/sceneCategories.html).

#### Earnings

Compensation for the prescan session was a show-up fee of $15 on top of their earnings from the prescan session. Compensation for the scan session was a show-up fee of $20 on top of their earnings from the scan session. Participants received an extra $50 for completing both sessions. Therefore, they could earn up to $145 for completing the entire study.

### Modeling the Value of (Instrumental) Information

Our general strategy was to implement linear mixed-effects regression to properly account for between-participant variance (fixed effects) and within-participant variance (random effects), using the MATLAB function fitlme with maximum likelihood estimation. In all mixed-effects models, we used the Satterthwaite approximation to calculate degrees of freedom, which has been shown to reduce Type 1 error compared to residual degrees of freedom (Luke 2017). Unless otherwise specified, each predictor was *z*-scored at the group level before the regression model was fit.

As a manipulation check, we first modeled participants’ bids *B* as a function of the probability distribution, **Figure 1A**), diagnosticity, and penalty *W* along with an intercept β_0_ observed variables on a trial: prior certainty (appears on screen as the maximum of the prior probability distribution, **Figure 1A**), diagnosticity, and penalty *W* along with an intercept *β*_0_ (**Equation 14**, Base Model). To account for the potentially confounding effect of slider movement on participants’ submissions, we included initial slider position as a nuisance regressors. We used fixed-effects terms for each variable and included random-effects terms for each variable by participant (**Equation 14**).

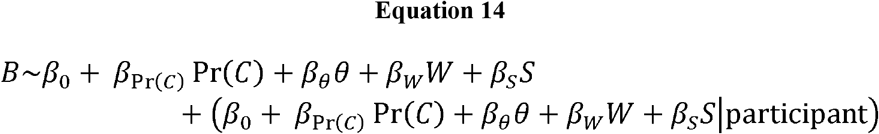

To model bid in terms of expected information gain (EIG), we developed a *condensed model* in terms of probability gain and an *extended model* decomposing probability gain into its mathematical components of prior certainty and expected posterior certainty. To model bid in terms of probability gain, we included fixed-effects terms for the intercept, probability gain, penalty, the interaction between probability gain and penalty (as suggested by the product of probability gain and penalty in **Equation 13**), and initial slider position, along with the corresponding random-effects terms by participant (**Equation 15**).

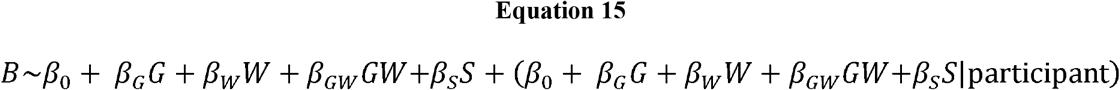

For the extended model, we replaced probability gain with its mathematical components (**Equation 16**), accounting for the possibility that a participant would not weight prior certainty and expected posterior certainty with equal magnitude when estimating the EIG of a sample.

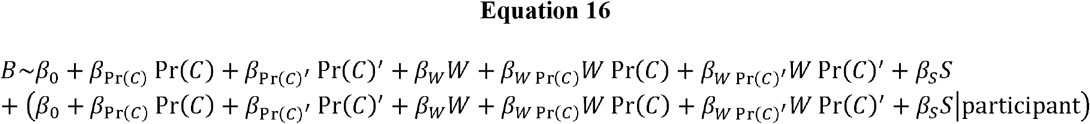

When we compared each participant’s coefficient on prior certainty to the coefficient on expected posterior certainty, we fit the extended model without *z*-scoring the predictors so that any difference between the coefficients was not attributable to a difference in the predictors’ standard deviations.

### fMRI Data Acquisition and Preprocessing

#### Acquisition

Whole-brain fMRI data were acquired on a 3-T Siemens MAGNETOM Prisma scanner with a 64-channel head coil at the Magnetic Resonance Imaging Center at the Zuckerman Mind Brain Behavior Institute of Columbia University. Functional images were acquired with a T2*-weighted, two-dimensional gradient echo spiral in/out pulse sequence (repetition time (TR) = 1,000 ms; echo time = 30 ms; flip angle = 52°, field of view = 230 mm; 2.4×2.4×2.4 mm voxels; 56 slices; multiband factor = 4). To reduce dropout in central frontal regions, slices were tilted by 10° forward from the AC-PC axis. During the scan session, the behavioral tasks were projected onto a mirror attached to the scanner head coil for the participant to see (Hyperion MRI Digital Projection System); participants made responses with the right hand through an MRI-compatible trackball (Current Design).

#### Preprocessing

Preprocessing was performed using the *fMRIPrep* pipeline, Version 1.5.0rc1 (Esteban et al. 2019) (RRID:SCR_016216). fMRIPrep uses a combination of tools from well-known software packages, including FSL, ANTs, FreeSurfer, and AFNI, and is based on *Nipype* 1.2.0 (Gorgolewski et al. 2011) (RRID:SCR_002502). For more details of the pipeline, see the section corresponding to workflows in *fMRIPrep*’s documentation at (https://fmriprep.org/en/latest/workflows.html).

### Anatomical data

The T1-weighted (T1w) image was corrected for intensity nonuniformity with N4BiasFieldCorrection (Tustison et al. 2010), distributed with ANTs 2.2.0 (Avants et al. 2008) (RRID:SCR_004757). The T1w image was then skull-stripped with a *Nipype* implementation of the antsBrainExtraction.sh workflow (from ANTs), using OASIS30ANTs as target template. Brain tissue segmentation of cerebrospinal fluid, white matter, and gray matter was performed on the brain-extracted T1w using fast (Zhang, Brady, and Smith 2001) (FSL 5.0.9, RRID:SCR_002823). Volume-based spatial normalization to Montreal Neurological Institute (MNI) space (MNI152NLin2009cAsym) was performed through nonlinear registration with antsRegistration (ANTs 2.2.0) (Fonov et al. 2009) (RRID:SCR_008796).

### Functional data

A skull-stripped susceptibility distortion–corrected BOLD reference was generated using a custom methodology of *fMRIPrep*. The BOLD reference was co-registered to the T1w reference using bbregister (FreeSurfer), which implements boundary-based registration using six degrees of freedom (Greve and Fischl 2009). Head-motion parameters (*x, y, z*, pitch, roll, and yaw) with respect to the BOLD reference were estimated before spatiotemporal filtering using mcflirt (FSL 5.0.9) (Jenkinson et al. 2002). BOLD runs were slice-time corrected using 3dTshift from AFNI 20160207 (Cox and Hyde 1997) (RRID:SCR_005927).

### fMRI Data Acquisition and Preprocessing

We used multi-voxel pattern analysis (MVPA) methods to identify regions in which the task variables were decodable. To do so, first, we used a whole-brain univariate general linear model (GLM) to estimate BOLD activation patterns (betas/parameter estimates) associated with each task variable. Then, we trained and tested a support vector regression decoder on the voxel-wise activation patterns that had been identified by the GLM. Univariate analyses were conducted using the GLM framework implemented in SPM12, Version 7487 (https://www.fil.ion.ucl.ac.uk/spm), convolving boxcar functions within the GLM by the SPM canonical hemodynamic response function. MVPA analyses were conducted using The Decoding Toolbox (Hebart, Görgen, and Haynes 2015; Görgen and Hebart 2022). Whole-brain statistical maps from functional data were overlaid on an average of the 23 participants’ individual T1-weighted (T1w) maps normalized to Montreal Neurological Institute (MNI) space. Since scanning did not occur during the Payout Stage, fMRI activation was only measured during the Bidding Stage.

*Whole-Brain Analyses to Localize Clusters in Which a Task Variable Was Decodable* Functional images normalized to MNI space were smoothed with a Gaussian kernel with a FWHM of 5×5×5 mm. Then, a condensed and an extended GLM was estimated for every participant from this normalized, smoothed functional time series. Both GLMs used a variable-epoch model (Grinband et al. 2008) using boxcar functions to represent each condition for each task variable during the decision period (the period between the beginning of the response window and the reaction time on trials that received a response, **Figure 1A**). The condensed GLM contained conditions for probability gain, penalty, bid, and slider displacement (i.e., the difference between the initial slider position and the slider position when the bid was submitted). The extended GLM replaced the probability gain conditions with conditions for prior certainty and expected posterior certainty. We fit these GLMs separately because probability gain is collinear with expected posterior certainty and prior certainty (because probability gain is the difference between the latter variables). The conditions for each variable were as follows, yielding one parameter estimate per run (four per participant):

- Probability gain: 0, 0.01–0.08, 0.09–0.17, 0.18–0.26, 0.27–0.35, 0.36–0.43 (nearly equally spaced bins on the range of probability gains with a separate category for 0, which was overrepresented)
- Prior certainty: low (0.5–0.53), medium (0.57–0.63), and high (0.87–0.93) (one for each level of prior certainty, “Selection of parameters”)
- Expected posterior certainty: low (0.57–0.63), medium (0.77–0.83), and high (0.87–0.93) (one for each level of expected posterior certainty)
- Penalty: $10 and $20 (one for each penalty condition)
- Bid: the bids discretized into 10 equally spaced bins over the available range ($0 to $9.88)
- Slider displacement: the signed slider displacements discretized into 10 equally spaced bins over the range of displacements during the scan session across all the participants

If the participant failed to respond to at least one trial during a run of the Bidding Stage, an additional boxcar function was added to the GLM to model the entire response window for each trial that they omitted. Finally, both GLMs also contained fixed-body motion-realignment regressors (*x, y, z*, pitch, roll, and yaw) and their respective first derivatives.

In the next step, a decoding analysis was performed on the parameter estimates of the GLM for each participant. A support vector regression was applied to each task variable of interest, trained and tested on all the variable’s conditions (across all the runs). Decoders were cross-validated using leave-one-run-out (four-fold) cross-validation. The label for each condition was the median value of the variable for the condition within the participant. The support vector regression was trained and tested on the same variable using a searchlight approach with a sphere of standard radius of 3 voxels (example: Kahnt et al. (2014)). Decoding accuracy for each voxel was measured as the Fisher’s *z*-transformed correlation coefficient between the decoder’s prediction and the true label for the variable. The searchlight analysis for probability gain used the condensed GLM. The searchlight analyses for prior certainty, expected posterior certainty, penalty, bid, and slider displacement used the extended GLM. Searchlight results were broadly similar for penalty, bid, and slider displacement between the two GLMs.

Finally, we identified significant clusters in which each variable of interest was decodable by submitting each participant’s accuracy map (across all brain voxels) for a variable to a second-level *T*-test, applying a cluster-wise correction for multiple comparisons using non-parametric permutation tests in SnPM13.1.08 (http://nisox.org/Software/SnPM13/) (Nichols and Holmes 2002), which have been shown to be most robust to false positives (Eklund, Nichols, and Knutsson 2016; Nichols et al. 2017). Permutation tests were based on a stringent cluster-forming height threshold of *p* < 0.001 and considered significant at a cluster-wise familywise error rate threshold of *p* < 0.05; we used 10,000 permutations (Holmes et al. 1996; Nichols and Holmes 2002).

#### Region of Interest (ROI) Analyses

Region-of-interest (ROI) decoding analyses were conducted on each significant cluster. ROI decoding was conducted the same way as the whole-brain decoding, except that the decoders were trained on all the voxels within a cluster instead of a searchlight sphere, yielding one accuracy statistic per ROI. This was done whether the decoder was trained and tested on the same variable or trained on one variable and tested on another (cross-decoding). All ROI decoding, including cross-decoding, was done with leave-one-run-out cross-validation. In all the expected posterior certainty (EPC) clusters, there was “substantial” evidence supporting the null hypothesis that cross-decoding accuracy of slider displacement from bid was not different from chance (BF > 10^1/2^).

## Acknowledgements

We would like to thank Chuen-Shin (Jessica) Cheng, Dania Elder, and Laura Hunter for help with data collection; Greg Jensen for help with data analysis; Kenneth Wengler for help with fMRI scanner sequences; and Mariam Aly, Brandon Ashinoff, Xiaosi Gu, Kenneth Wengler, Daniel Wolpert, and Michael Woodford for helpful advice on the project and manuscript. This study was funded by the Seed Grant for MR Studies Program of the Zuckerman Mind Brain Behavior Institute at Columbia University (CU-ZI-MR-S-0011) and the National Science Foundation Graduate Research Fellowship Program (DGE-1644869, NMS). This work was also supported by the National Institute of Mental Health under awards R01MH117323 and R01MH114965 (GH).

## Footnotes

1 Letting be the agent’s bid for the sample, the probability of receiving a sample is therefore 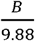 because Pr(*X<9*.*88=0Bfxdx=0B19*.*88dx=B9*.*88*.

## Notes

### Competing Interest Statement

The authors have declared no competing interest.

