## Supplement for "A Distinct Neural Code Supports Prospection of Future Probabilities During Instrumental Information-Seeking"

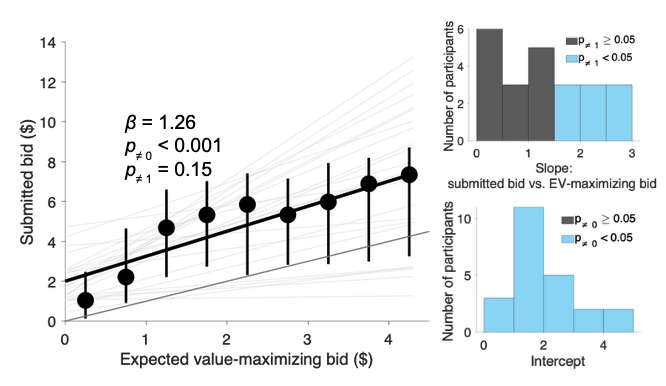

**Figure S1.** Across trials in the scan session, participants in the Main Cohort (*N* = 23) submitted bids (*y*-axis) that were positively and monotonically associated with the trials’ corresponding expected value–maximizing bids (*x*-axis): the fixed-effects (group-level) regression coefficient (slope) was significantly greater than 0 (fixed-effects regression coefficient = 1.26, SE = 0.17, *T*(22.99) = 7.37, *p*_≠ 0_ < 0.001). The coefficient was also not significantly different from unity, suggesting lack of evidence that bids increased at a different rate with VOI from the expected value–maximizing bid (fixed-effects regression coefficient = 1.26, SE = 0.17, *T*(22.99) = 1.50, *p*_≠ 1_ = 0.15). The distribution of individual participants’ slopes is in the upper histogram; most participants’ slopes (14 of 23) were *not* significantly different from unity.

However, the fixed-effects intercept was significantly greater than 0 (fixed-effects intercept = 2.00, SE = 0.24, *T*(23.01) = 8.30, *p*_≠0_ < 0.001), indicating that participants generally bid more than the expected value–maximizing bid. The distribution of individual participants’ intercepts is in the lower histogram; all participants’ intercepts were significantly greater than 0.

The black diagonal line is the least-squares regression line for the group while the thin, gray lines are the least-squares lines for each participant. For visualization, group submitted bids are binned by expected value–maximizing bid; however, the regression was carried out on the raw bids, not the binned medians. Error bars represent interquartile range. The long, thick gray diagonal represents unity.

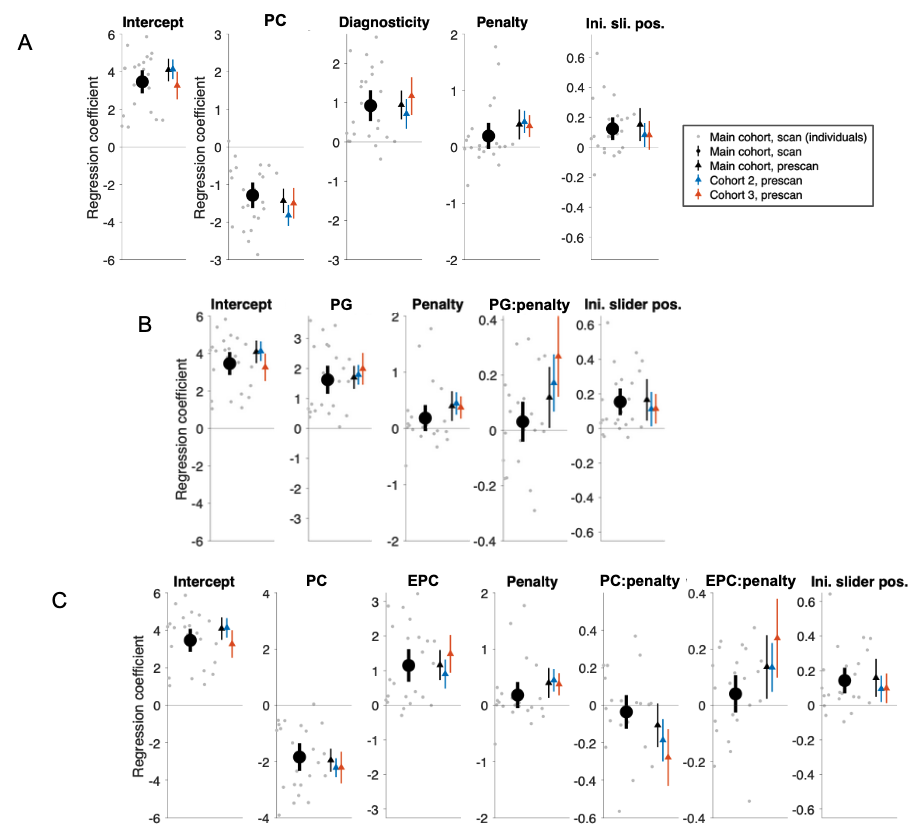

**Figure S2.** **Full set of regression coefficients from the behavioral models**. Coefficients from the Base (**A**), Condensed (**B**), and Extended (**C**) models are shown in the same format as **Figure 2A–C**. The main coefficients for diagnosticity, PC, penalty, PG, and EPC are replicated from **Figure 2**, and shown alongside all the other coefficients in each model. Because regressors were z-scored before the model was fit, the intercepts estimate the predicted bid when the regressors were at their means. Initial slider position had a positive but negligible effect on bid compared to the other variables, as did the interaction between penalty and probabilistic quantities.

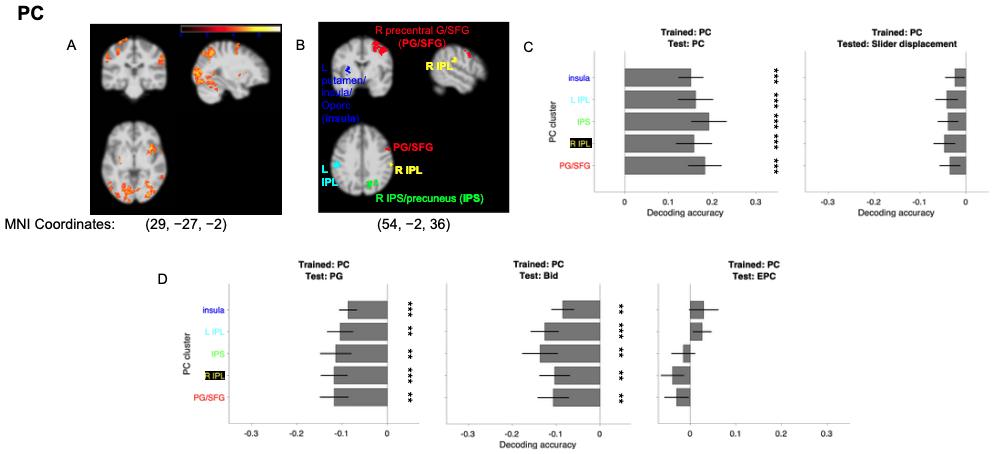

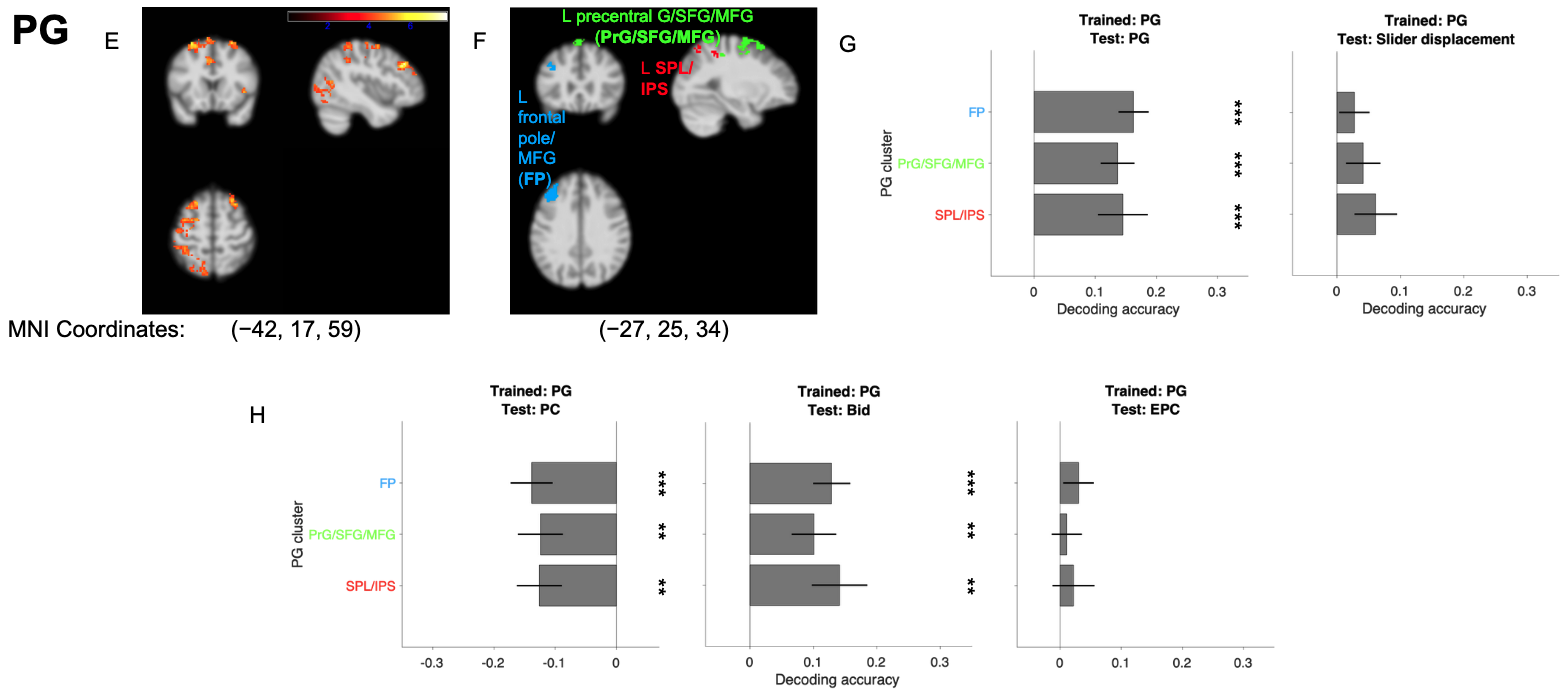

**Figure S3.** **Clusters identified by a whole-brain searchlight as showing significant decoding of PC and PG.** The format for all panels is the same as in **Figure 3A–B: Clusters showing decoding of PC** were identified in the left putamen/insula/operculum, (dark blue), left inferior parietal lobule (IPL, cyan), right intraparietal sulcus (IPS)/precuneus (green), right IPL (yellow) and right precentral gyrus/superior frontal gyrus (SFG, red). **C:** The PC clusters do not show significant cross-decoding with slider displacement (right), **D:** The PC clusters show significant cross decoding of PG and bid (left and middle panels; the negative values indicate the negative association of these quantities with PC), but do not significantly cross-decode EPC (right panel). **E–F:** **Clusters showing decoding of PG** were identified in the left frontal pole/middle frontal gyrus (MFG; light blue) left precentral gyrus/superior frontal gyrus (SFG/MFG, green) and left superior parietal lobule (SPL/IPL, red). **G:** The PG clusters do not show significant cross-decoding with slider displacement (right). **H:** The PG clusters show significant cross decoding of PG and bid (left and middle panels; the negative values indicate the negative association between PG and PC), but do not significantly cross-decode EPC (right panel).

**Table S1.** Peak voxels from the three significant clusters in which expected posterior certainty (EPC) was decodable (**Figure 3B**; cluster-forming height threshold: *p* < 0.001; cluster-wise family-wise error rate correction: 0.05). Anatomical regions were assigned by the JuBrain/SPM Anatomy Toolbox 3.0 (Eickhoff et al. 2005; 2006; 2007).

| Cluster | Size (vox.) | *T*-statistic | MNI Coordinates | | | Anatomical region |
| --- | --- | --- | --- | --- | --- | --- |
|  |  |  | *x* | *y* | *z* |  |
| Right intraparietal sulcus/extrastriate cortex (IPS) | 270 | 6.19 | 29 | −84 | 25 | hIP7 (IPS) |
|  |  | 5.79 | 7 | −86 | 20 | hOc2 [V2] |
|  |  | 5.47 | 26 | −67 | 32 | hIP8 (IPS) |
|  |  | 5.12 | 33 | −79 | 25 | hIP4 (IPS) |
|  |  | 5.03 | 29 | −82 | 35 | hPO1 (IPS) |
|  |  | 4.92 | 24 | −94 | 28 | hIP7 (IPS) |
|  |  | 4.75 | 12 | −79 | 23 | hOc3d [V3d] |
|  |  | 4.59 | 33 | −84 | 35 | hIP7 (IPS) |
|  |  | 4.52 | 7 | −82 | 18 | hOC2 [V2] |
|  |  | 4.50 | 14 | −82 | 32 | hPO1 (IPS) |
| Right occipital fusiform gyrus/cerebellum (OFG) | 163 | 6.94 | 31 | −63 | −11 | FG1 |
|  |  | 4.91 | 17 | −65 | −16 | hOC3v [V3v] |
|  |  | 4.89 | 26 | −65 | −13 | FG1 |
|  |  | 4.87 | 12 | −74 | −11 | hOc3v [V3v] |
|  |  | 4.75 | 19 | −70 | −16 | hOc3v [V3v] |
|  |  | 4.58 | 21 | −67 | −13 | hOc4v [V4(v)] |
|  |  | 4.14 | 19 | −79 | −8 | hOc3v [V3v] |
|  |  | 4.13 | 24 | −72 | −11 | hOc4v [V4(v)] |
|  |  | 4.01 | 14 | −79 | −8 | hOc3v [V3v] |
|  |  | 3.82 | 29 | −77 | −8 | hOc4v {V4(v)] |
| Right occipital pole (OP) | 105 | 5.98 | 5 | −98 | 4 | N/A |
|  |  | 5.49 | 5 | −94 | −6 | hOc1 [V1] |
|  |  | 5.01 | 12 | −101 | 4 | hOc1 [V1] |
|  |  | 4.61 | 9 | −101 | 11 | hOc2 [V2] |
|  |  | 4.42 | 12 | −94 | −11 | hOc2 [V2] |
|  |  | 4.07 | 17 | −91 | −13 | hOc3v [V3v] |
|  |  | 4.02 | 14 | −101 | −11 | hOc1 [V1] |
|  |  | 3.89 | 9 | −96 | 8 | hOc1 [V1] |
|  |  | 3.68 | 9 | −91 | −13 | hOc2 [V2] |
